# Effects of protoscoleces excretory-secretory products of *Echinococcus granulosus* on hepatocyte growth, function, and glucose metabolism

**DOI:** 10.1101/2022.10.04.510778

**Authors:** Guangyi Luo, Haiwen Li, Qiong Lu, Jiangtao Cao, Hailong Lv, Yufeng Jiang

**Affiliations:** School of Medicine, Southwest Jiaotong University, The Affiliated Hospital of Southwest Jiaotong University, Medical Research Center, The Third People’s Hospital of Chengdu, Chengdu, Sichuan, China; Section for Hepatopancreatobiliary Surgery, Department of General Surgery, The Third People’s Hospital of Chengdu, Affiliated Hospital of Southwest Jiaotong University, Chengdu, Sichuan, China; Department of Hepatobiliary Surgery, Anyue County People’s Hospital, Ziyang, Sichuan, China; School of Basic Medicine, Chengdu Medical College, Chengdu, Sichuan, China; Department of Infectious Diseases, Anyue County People’s Hospital, Ziyang, Sichuan, China

**Author notes:** **Corresponding author**, (HL), (YJ).

**Keywords:** *Echinococcus granulosus*, excretory-secretory products, hepatocyte, cellular function, glucose metabolism

## Abstract

Cystic echinococcosis (CE) is one of the most widespread and harmful zoonotic parasitic diseases and it most commonly affects the liver. In this study, we characterized multiple changes in mouse hepatocytes following treatment with excretory-secretory (ES) products of *Echinococcus granulosus* protoscoleces by a factorial experiment. The cell counting kit-8 assay (CCK-8), the 5-ethynyl-2’-deoxyuridine (EdU) assay, and flow cytometry were used to detect the growth of hepatocytes. Inverted microscopy, scanning electron microscopy (SEM), and transmission electron microscopy (TEM) were used to observe the morphology and ultrastructure of hepatocytes. An automatic biochemical analyzer and an ELISA detection kit were used to determine six conventional hepatocyte enzymatic indices, the levels of five hepatocyte-synthesized substances, and the contents of glucose and lactate. Western blot analysis was conducted to analyze the protein expression of six rate-limiting enzymes in the glucose metabolism pathway in hepatocytes: glutamic-pyruvic transaminase (ALT), glutamic-oxalacetic transaminase (AST), alkaline phosphatase (ALP), lactate dehydrogenase (LDH), gamma-glutamyl transpeptidase (GGT), and leucine arylamidase (LAP). The results of the CCK-8 and EdU assays both showed that ES could inhibit the proliferation of hepatocytes, and flow cytometry indicated that ES could promote apoptosis of hepatocytes. After ES treatment, the ultrastructure of hepatocytes was disrupted to a certain extent. The changes in the cell membrane and microvilli were observed through SEM, and the changes in the nucleus, mitochondria, and rough endoplasmic reticulum were observed through TEM. After ES treatment, the enzymatic activities of the six hepatocyte enzymes were increased in addition to the Fe metabolism and the synthesis of albumin (ALB), uric acid (UA), and urea, whereas the synthesis of transferrin (TRF) was decreased. The expression levels of all six key enzymes in the glucose metabolism pathway in hepatocytes were decreased, and the biological effects were significantly inhibited. We analyzed the causes and possible complications caused by various changes and advocate corresponding measures. We also propose possible mechanisms by which protoscoleces cause hepatocyte necrosis, but the specific mechanism requires further study.

**Author Summary:** *Echinococcus granulosus*, the most widely distributed and most infected tapeworm, has caused serious economic and social burdens to pastoral areas in China. The metacestodes of *Echinococcus granulosus* mainly infect the liver of intermediate hosts (humans, cattle, sheep, etc.). Currently, the effects of *Echinococcus granulosus* on hepatocytes’ ultrastructure, enzymology, function, and glucose metabolism have not been characterized, and accurate characterization is crucial in the study of related pathogenesis and preventive therapy. Here, we characterize multiple changes in hepatocytes using excretory-secretory (ES) products of *Echinococcus granulosus* protoscoleces and mouse hepatocyte action by a factorial experiment. We found that ES inhibited hepatocyte proliferation and promoted hepatocyte apoptosis. ES can cause a certain degree of damage to the cell membrane, nucleus, mitochondria, and endoplasmic reticulum of hepatocytes. After ES treatment, six enzymatic indexes of hepatocytes were elevated, they were ALT, AST, LDH, ALP, GGT, and LAP, and Fe, ALB, UA, and urea levels synthesized by hepatocytes were significantly higher and TRF levels were significantly lower. Reduced expression of rate-limiting enzymes in six pathways of glucose metabolism in hepatocytes, including PFK-1, IDH, G-6-PD, GS, GP, and GLUT-2, indicating that ES inhibits glucose metabolism in hepatocytes. Our study not only characterized the effects of ES on hepatocytes in detail but also proposed the possible mechanisms causing these effects, which provided a basis for subsequent studies on related pathogenesis and prevention, and treatment.

## Introduction

Cystic echinococcosis (CE) is a zoonotic parasitic disease caused by metacestodes of *Echinococcus granulosus (E. granulosus*) parasitizing in suitable intermediate hosts[1]. CE has been listed as one of the 17 Neglected Tropical Diseases by the World Health Organization[2]. It is worth mentioning that western China is one of the countries with high CE prevalence, which causes serious economic and social losses in the endemic areas[3]. Intermediate hosts (such as sheep and cattle) are occasionally infected by swallowing *E. granulosus* eggs, which hatch into oncospheres in the duodenum and then pass through the intestinal mucosa into the portal vein system. Most of the oncospheres are blocked in the liver, but a few can flow to the lungs through the hepatic sinusoid and even form lesions in the brain and other parts of the body. Totally, the liver is the organ that is most affected by hydatid cysts[4]. Oncospheres develop firstly into small cysts in the liver, which grow large and crush the liver parenchyma, forming a cystic mass (hydatid cyst) with a multilayered wall structure. The cyst wall of hydatid cysts is divided into two layers: (1) the inner capsule, which has a worm structure; and (2) the outer capsule, which is a dense fibrous layer characterized by macrophage granulomatous lesions and fibrosis formed by the host upon immune rejection of the parasite[5]. At present, the treatment strategy for CE is surgical resection and drug therapy, but surgical resection can result in secondary infection and drug resistance is gradually increasing[6–9].

Followed by hydatid cysts gradually develop and mature, protoscoleces (PSCs) emerge in the cyst and become new sources of infection. As multicellular organisms, PSCs and their hosts rely on complex cellular signaling mechanisms to properly regulate development, homeostasis, and physiological functions. The mechanism is that certain cell types release peptides or lipophilic hormones and cytokines, and receptor molecules on target cells or tissues induce appropriate cellular responses [10]. Proteomic analysis showed that *E. granulosus* PSCs excrete or secrete specific macromolecular proteins to act on the host and regulate the expression of host genes[11]. As hydatid cysts grow, they compress the surrounding tissues and organs, causing cells and tissues to atrophy and inducing necrosis[12]. This process might be mediated by the immune response, apoptosis, and Toll-like receptors[13–17]. Immunohistochemical studies showed that iNOS, TNF-α, NF-κB, Vimentin, Bcl-2, and CD68 levels were significantly increased in liver tissue affected by hydatid cysts, which was correlated with the abundance of collagen and reticulin fibers[18]. Another study revealed that both iNOS and IL-10 macrophage types were involved in the local immune response to hydatid cysts and that Th1 and Th2 immune response stimuli were sustained simultaneously[19]. Additionally, PSCs coulde promote the protein expression of hepatocyte keratin, caspase-3, caspase-9, and Bax, as well as reduce the expression of Bcl-2, inhibit the proliferation of hepatocytes, and promote hepatocyte apoptosis. Moreover, the mRNA expression of CK8 and CK18 in tissues affected by *Echinococcus* spp. is higher than that in normal tissues, which reflects the liver injury caused by CE in vitro[20].

Hydatid cysts parasitizing their host as metacestodes interact with the host at many levels. The PSCs that sprout in hydatids are in their original form and are the main cause of disease development during host infection. The excretory-secretory (ES) products of hydatid cysts, which are involved in a series of biochemical reactions, are the main agents mediating the interaction between hydatid cyst and host. At present, it is not known how hydatid cysts cause hepatocyte degeneration, atrophy, and necrosis or how the outer capsule is formed. Moreover, little is known about the factors affecting innate susceptibility and resistance to *E. granulosus* infection. Therefore, understanding the effects of hydatid cyst growth and development on host hepatocytes is of great significance for exploring the pathogenesis, treatment, and prevention of CE. The purpose of this study was to explore the effect of hydatid cysts on hepatocyte growth and function by means of the interactions between ES of *E.granulosus* protoscoleces and mouse hepatocytes. We provided the basis for exploring the molecular mechanism of hepatocyte apoptosis, necrosis and disappearance caused by hydatid cysts during growth and development of intermediate hosts.

## Materials and Methods

### In vitro culture and ES product collection of *E. granulosus* PSCs

Livers from sheep naturally infected with *E. granulosus* PSCs were collected from the Shihezi slaughterhouse, Xinjiang, China (Fig. 1A). The cyst fluid containing *E. granulosus* PSCs was aspirated under aseptic conditions, and the inner capsule was washed several times to isolate the PSCs. The PSCs were collected in a sterile centrifuge tube, filtered twice through a sterile stainless-steel filter with a pore size of 200 μm, and rinsed 5—10 times with PBS containing a double antibiotics until the PSCs naturally sank in the suspension and the liquid was clear. The treated PSCs were stained with 0.4% Trypan Blue to determine their viability. The PSCs with viability greater than 95% were used in this study, and the optimal culture density of PSCs (2000 PSCs/ml) was cultured as recommended previously[21]. This density is consistent with our calculation of a density of 400,000 PSCs in 200 ml of cyst fluid in a single hydatid cyst. PSCs were cultured in DMEM medium (Hyclone, USA) containing 10% fetal bovine serum (Every green, China), 100 U/ml penicillin, and 100 μg/ml streptomycin (Hyclone, USA) at 37°C and 5% CO_2_. During the first three days of culture, the culture medium was changed daily to remove host impurities, and purified PSCs were obtained (Fig. 1B). The culture supernatant was collected 48 h after each subsequent culture change[11], which contained ES products of the PSCs. The ES products were sterilized through a 0.22-μm filter to obtain pure ES products (100% ES) for subsequent experiments. We did not concentrate the ES products. Normal NCTC-1469 hepatocytes were purchased from Procell biotechnology company (China).

**Fig 1.**
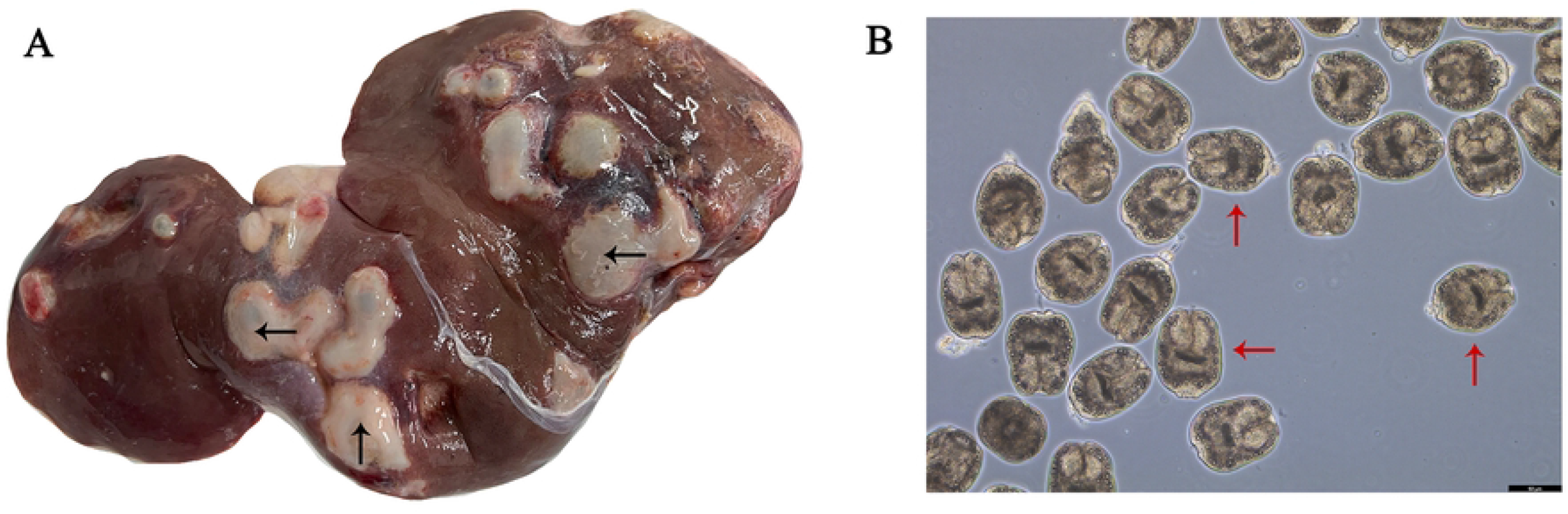
Liver and PSCs. (A) Liver of sheep infected with *E. granulosus* contains hydatid cysts. (B) Purified PSCs (×200). Hydatid cysts and PSCs are marked with black and red arrows, respectively.

### Cell counting kit-8 (CCK-8) assay

We first used the cell counting kit-8 (CCK-8) assay (Biosharp, China) to determine the viability of hepatocytes. NCTC-1469 cells in the logarithmic growth phase were counted and the cell density was adjusted to 5×10^4^/ml. Next, 100 μl of cell suspension per well was seeded in 96-well plates. The cultures were incubated with different concentrations of ES (0%, 10%, 25%, 50%, 75%, and 100%) for 24, 48, 72, and 96 h, respectively. CCK-8 reagent was diluted at 1:10 in a serum-free medium, and one plate of cells was taken every day. After the culture medium was aspirated, 110 μl of diluted CCK-8 working solution was added to the well, and the culture was kept at 37°C and 5% CO_2_ for 0.5–1 h. The optical density (OD) of each well was measured at 450 nm using a microplate reader (Molecular Devices, America), and then the inhibition rate was calculated as follows: inhibition rate = [(Ac - As)/(Ac - Ab)] × 100%, where As is the OD of experimental wells, Ac is the OD of control wells (0% ES), and Ab is the OD of blank wells (no cells and 0% ES).

### 5-ethynyl-2’-deoxyuridine (EdU) assay

To verify the effect of ES treatment on the proliferation of NCTC-1469 cells, the proliferation of hepatocytes was also detected by immunofluorescence microscopy. A factorial experiment was designed, with ES and time as independent variables and the experimental result as a dependent variable. There were four groups in the experiment: control-24 h, control-48 h, ES-24 h, and ES-48 h. The EdU Cell Proliferation Kit with Alexa Fluor 488 (Beyotime, China) was used to determine the proliferation of hepatocytes. In brief, hepatocytes were incubated with a EdU working solution (10 μM) for 2 h at 37°C in darkness. Then the cells were fixed with methanol for 15 min at ambient temperature. Then the cells were permeated with 0.3% Triton X-100 for 15 min, incubated with Click solution for 30 min, washed, and stained with 1 ml Hoechst 33342 for 5 min. Finally, the images were captured under an immunofluorescence microscope (Olympus, Japan).

### Detection of hepatocyte apoptosis

We used an Annexin V-APC/PI Apoptosis Detection Kit (KeyGen Biotech, China) to detect hepatocyte apoptosis by flow cytometry. The cells were resuspended with 500 μl of Binding Buffer, then 5 μl of Annexin V-APC and 5 μl of PI were added, and the cells were exposed to light at room temperature for 15 min. The cells were observed and analyzed by a flow cytometry analyzer (Beckman, USA).

### Scanning electron microscopy and transmission electron microscopy

The ultrastructure of hepatocytes before and after ES treatment was analyzed by scanning electron microscopy (SEM) and transmission electron microscopy (TEM). The cells were observed before and after ES treatment under an inverted microscope. The cells were washed twice with PBS, fixed with 0.5% glutaraldehyde for 10 min, and again washed twice with PBS. Finally, 1 ml of 3% glutaraldehyde fixation solution was added and samples were stored at 4°C. For SEM, the cells were washed twice with ultrapure water for 5 min each time and dehydrated with a series of gradient alcohol (30%, 50%, 70%, 80%, 90%, 95%, and 100%) for 10 min per step. The slides were gently glued to the conductive adhesive and treated with ion sputtering spray (Japan Hitachi Nake High-Tech Enterprise). Finally, a suitable position was selected under a scanning electron microscope (FEI, USA) for observation in appropriate multiples. For TEM, the cells were prefixed with 3% glutaraldehyde, postfixed in 1% osmium tetroxide, treated with a diuretic in a series of acetone concentrations, infiltrated with Epox 812, and embedded. The semithin sections were stained with methylene blue. Ultrathin sections were cut with a diamond knife and stained with uranyl acetate and lead citrate. Sections were examined with a JEM-1400-FLASH Transmission Electron Microscope (JEOL, Japan).

### Detection of enzyme activities and synthesis function in hepatocytes

The culture supernatant was collected into centrifuge tubes, and the impurities were removed by a 0.22-μm filter. An automatic biochemical analyzer (HITACHI, Japan) was used to determine the activities of glutamic-pyruvic transaminase (ALT), glutamic-oxalacetic transaminase (AST), alkaline phosphatase (ALP), lactate dehydrogenase (LDH), gamma-glutamyl transpeptidase (GGT), and leucine arylamidase (LAP) to evaluate the effects of ES on hepatocyte injury. Albumin (ALB), transferrin (TRF), urea, uric acid (UA), and Fe were detected by an automatic biochemical analyzer and an ELISA kit (Hlkbio, China) to evaluate the effects of ES on the synthesis function of hepatocytes. In addition, the concentrations of glucose (Glu) and lactic acid (LA) in the supernatant were detected by ELISA.

### Western blot analysis

To investigate the effects of ES on glucose metabolism of hepatocytes, the expression levels of rate-limiting enzymes of various glucose metabolism pathways were determined by Western blot (WB) analysis. These enzymes included phosphofructokinase-1 (PFK-1) for anaerobic oxidation of glucose, isocitrate dehydrogenase (IDH) for aerobic oxidation of glucose, glucose-6-phosphate dehydrogenase (G-6-PD) for the pentose phosphate pathway, glycogen synthase (GS) for glycogen synthesis, glycogen phosphorylase (GP) for glycogen decomposition, and glucose transporter-2 (GLUT-2) for glucose transport. The above six enzymes are not only the main rate-limiting enzymes of these glucose metabolism pathways, but also the main regulatory factors of these processes. Hepatocytes were lysed with RIPA buffer and proteins were collected. Protein concentrations were determined with a BCA protein quantification kit. After extraction, the proteins were denatured by heating, separated by SDS-PAGE, and transferred to PVDF membranes, which were blocked in 5% skim milk diluted with TBST buffer for 2 h at room temperature. PVDF membranes were incubated overnight at 4°C with the following primary antibodies: rabbit anti-G-6-PD (1:5000, Proteintech, China), rabbit anti-GLUT-2 (1:1000, Proteintech, China), rabbit anti-GS (1:1000, Zen-bio, China), rabbit anti-IDH (1:2000, Proteintech, China), rabbit anti-PFK-1 (1:1000, Zen-bio, China), rabbit anti-GP (1:2000, Zen-bio, China), and rabbit anti-β-actin (1:50,000, ABclonal, China). Next, the membranes were washed, incubated for 2 h at room temperature with secondary antibody (1:5000), and washed three times with TBST. Protein bands were visualized using ECL luminescence solution.

### Statistical analysis

SPSS 28.0 software was used to perform all statistical analyses. All quantitative data are expressed as mean ± SD. Experiments were repeated at least three times unless otherwise stated. The Levene test was used to prove the homogeneity of variance. Two-way ANOVA was used for comparison between sample means. The simple effect was compared if there was an interaction, and the main effect was compared otherwise. We only focused on the main effects of ES. For the simple effect, we compared ES-24 h and control-24 h, ES-48 h and control-48 h, and ES-48 h and ES-24 h. Comparisons of other groups were not meaningful. In all statistical plots, the *P*-values for the interaction and ES main effects are replaced by P_i_ and P_E_, respectively, and Pη^2^_i_ and Pη^2^_E_ as partial η^2^ represent their corresponding effect sizes, respectively. *P*-values of < 0.05 were considered to indicate significant differences (\**P* < 0.05, \*\**P* < 0.01).

## Results

### Effects of ES on hepatocyte activity, proliferation, and apoptosis

ES inhibited the proliferation and induced apoptosis of NCTC-1469 cells. The CCK-8 assay showed that ES inhibited the proliferation of NTCC-1469 in a concentration- and time-dependent manner (Fig. 2A). In the first 48 h of ES treatment, the inhibitory effect of ES on the proliferation of NTCC-1469 cells was gradually enhanced, and the inhibitory effect was gradually weakened after 48 h. Higher ES concentrations were associated with stronger inhibitory effects on NCTC-1469 cells. The maximum inhibitory effect of ES was observed at 48 h. To reflect the effect of time, two time points (24 and 48 h) were selected for <u>exploration</u>. In immunofluorescence experiments, cell images were obtained before and after ES treatment (Fig. 2B). ImageJ software was used to analyze the images and calculate the proportion of EdU-positive cells, i.e., proliferating cells, in each group (Fig. 2C). We found an interaction between ES and time. The proportion of EdU-positive cells was significantly lower in the ES-48 h group than in the control-48 h group. Moreover, the proportion of EdU-positive cells was lower in the ES-48 h group than in the ES-24 h group. ES inhibited the proliferation of NCTC-1469 cells, and this inhibitory effect was gradually enhanced with the prolonging of time. To determine the effects of ES on apoptosis, flow cytometry analysis was conducted, and scatter plots were obtained (Fig. 3A). We found no interaction between ES and time. The apoptosis rate was significantly increased after the addition of ES, indicating that ES induced hepatocyte apoptosis (Fig. 3B).

**Fig 2.**
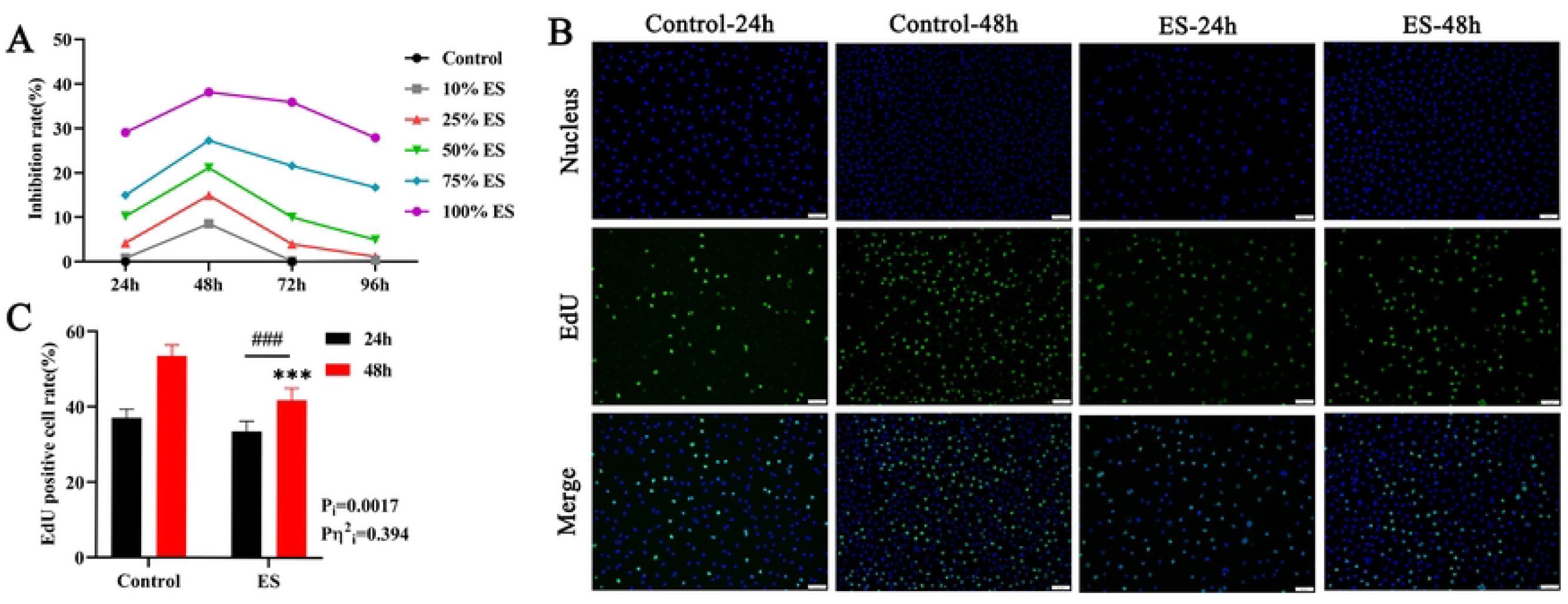
Effects of ES on hepatocyte proliferation. (A) The effects of different concentrations of ES on hepatocyte viability were detected by CCK-8. (B and C) Immunofluorescence images of positive (proliferating) cells (×200) (B) and EdU positive cell rates (C) after 24 and 48 h of ES treatment. Proliferating cells gave a green fluorescence signal, while the nuclei of all viable cells gave a blue fluorescence signal. *\*P* < 0.05 vs. control group, #*P* < 0.05.

**Fig 3.**
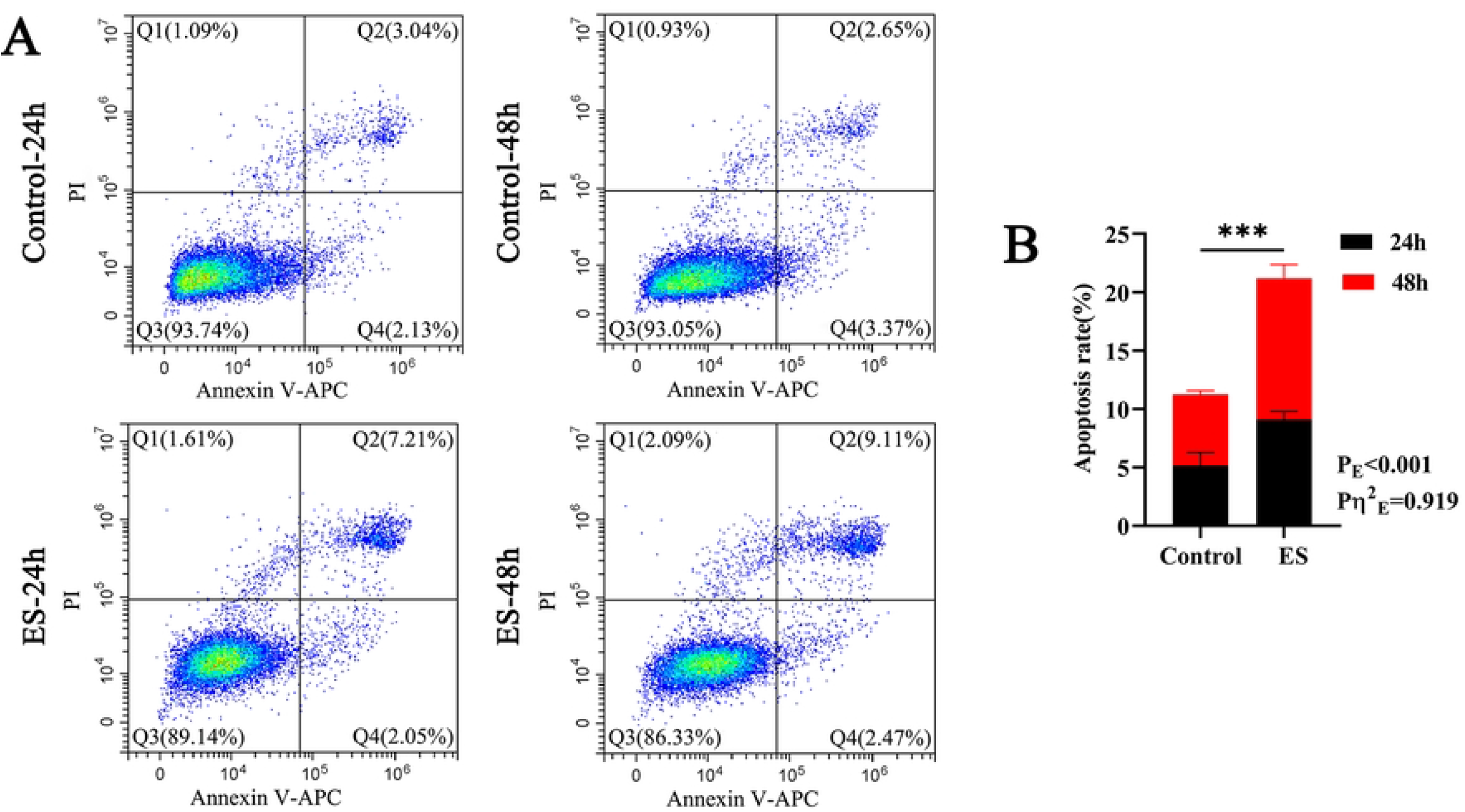
Apoptosis of hepatocytes after ES treatment. (A) Scatter plot showing the flow cytometry analysis results in each group after ES treatment. Q1 indicates dead cells or cell debris, Q2 indicates middle and late apoptotic cells, Q3 indicates viable cells, and Q4 indicates early apoptotic cells. (B) The apoptosis rate of each group is expressed as follows: apoptosis rate = Q2 + Q4. \**P* < 0.05.

### Effects of ES on ultrastructure of hepatocytes

Inverted microscope images were taken before and after ES treatment (Fig. 4A). The number of cells was higher in the ES group than in the control group, and there was no significant difference in cell morphology. In the control group, the cells were round or oval, few were polygonal, and the nuclei were round and large, located in the center of the cells. Most cells in the ES group were polygonal or spindle-shaped, while a few were round or oval. Some of the nuclei in the ES group showed irregular shapes and different positions. Compared with the control group at the same time point, we mainly found that the cell morphology changed from round or oval to polygonal or spindle shaped.

**Fig 4.**
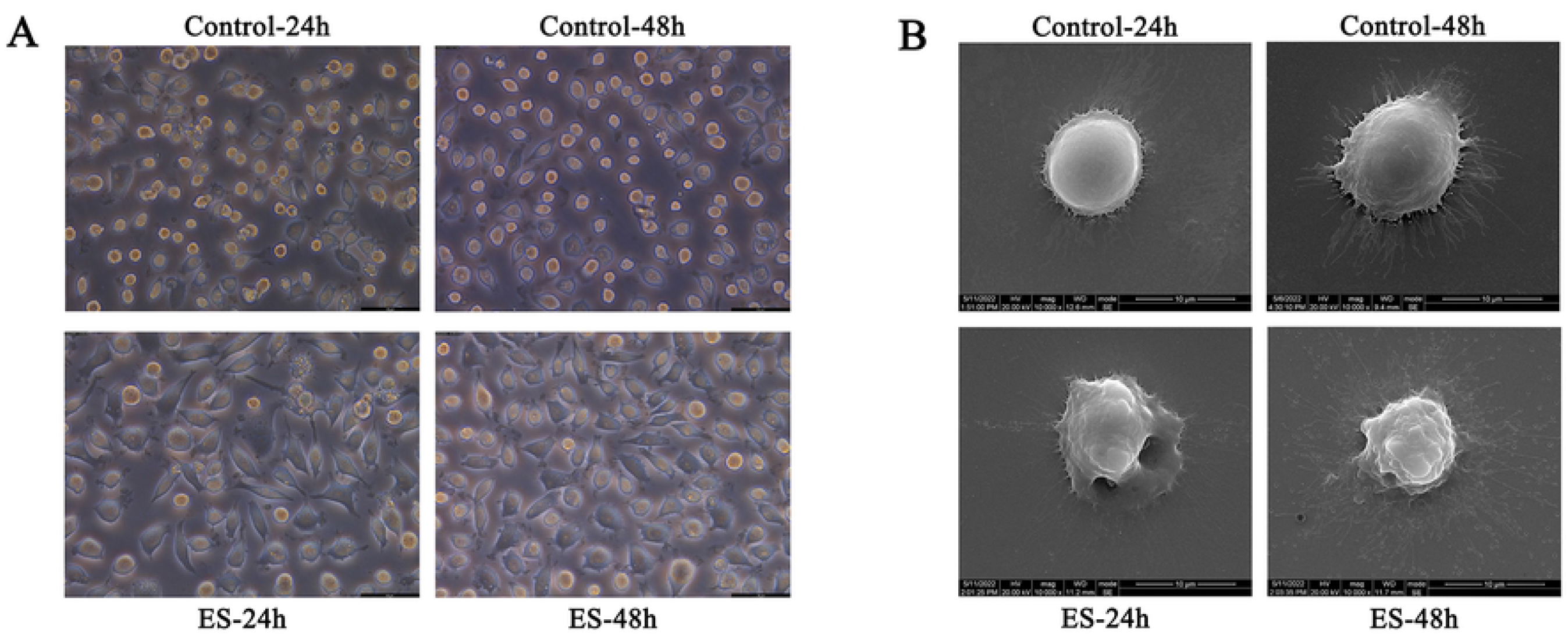
Morphological and structural changes of hepatocytes after ES treatment. Cells were observed under an inverted microscope (×200) (A) and a scanning electron microscope (×10,000) (B).

SEM analysis (Fig. 4B) revealed that the cells in the control-24 h group were regular in shape, round or oval, with intact cell membranes and a clear microvillous structure. Most cells in the control-48 h group showed regular morphology, and the number of microvilli was higher than that in the control-24 h group. In the ES-24 h group, the cells were slightly deformed and smaller, the cell membrane was intact but appeared foaming and depressed, fewer microvilli were present, and the cells were visibly damaged. In the ES-48 h group, the cell morphology was extremely irregular, the cell structure collapsed, the cell membrane was disrupted, surface foaming and depression were prominent, the microvilli fell off, and the cells seemed to die.

TEM analysis revealed the structural changes of organelles such as the nucleus (N), mitochondria (Mit), and rough endoplasmic reticulum (RER) (Fig. 5). Not all organelles could be observed. In the control-24 h group, the cell structure was normal, the nucleus was oval, the chromatin (CH) was evenly distributed, the nucleolus (NUE) was clearly visible, and the nuclear envelope (NE) was continuous and intact. Mit was abundant in the cytoplasm, with a short rod-like and clear structure, scattered or distributed in piles in the cytoplasm. The RER was linear, without vacuole-like structure and endoplasmic reticulum expansion. It was difficult to identify intracellular organelles such as the Golgi apparatus, the smooth endoplasmic reticulum, ribosomes, and lysosomes. In the control-48 h group, the cell morphology and structure were normal and organelles such as Mit and RER were intact and clear. No large differences were observed between the control-48 h and control-24 h groups. In the ES-24 h group, the CH of some cells lacked a diffuse distribution and was gradually condensed into clumps, and the NE began to fold. Some Mit was slightly swollen, and in some cells, the RER was slightly dilated. In the ES-48 h group, nuclear morphology was significantly changed, with some nuclei being pyknotic, the CH edges converged, and the NE surface was rough. The number and volume of Mit in the cytoplasm were significantly decreased, and in some cells, the RER was significantly expanded.

**Fig 5.**
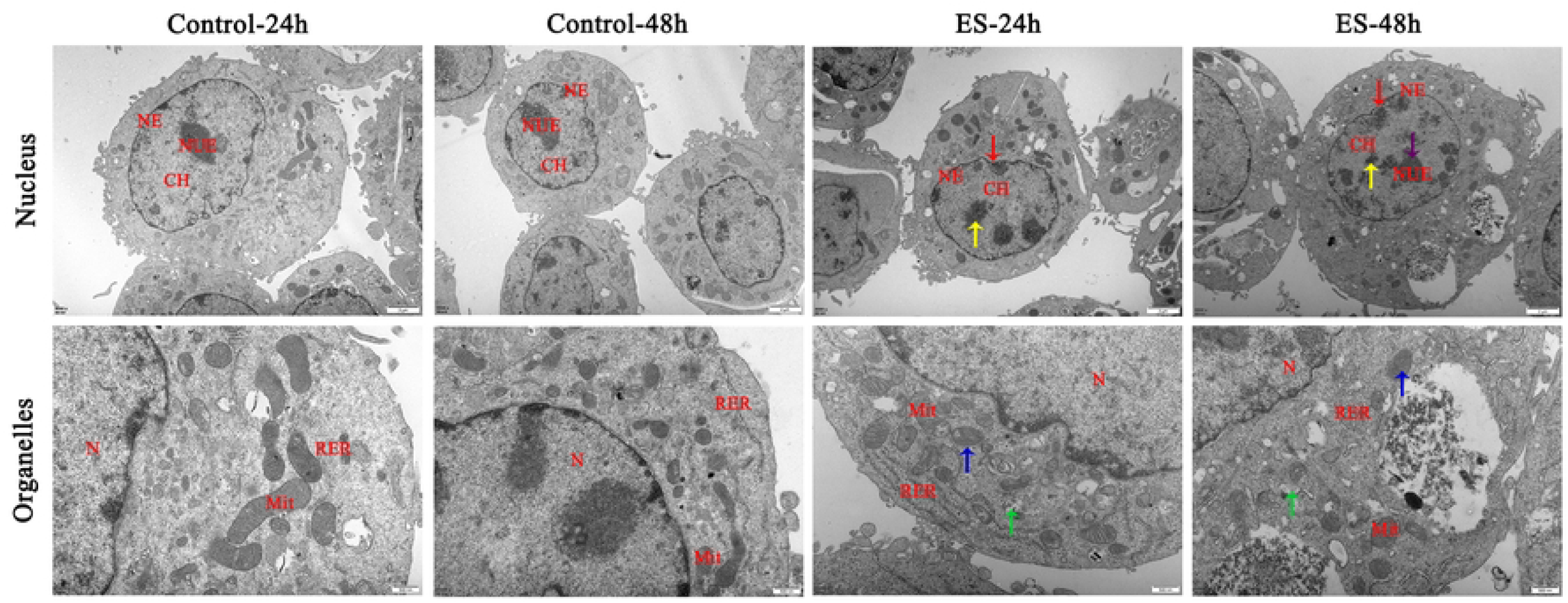
Ultrastructural changes of hepatocytes after ES treatment. In the nucleus (×8000), chromatin (CH), nucleolus (NUE), and nuclear envelope (NE) are indicated by yellow, purple, and red arrows, respectively. In the organelles (×25,000), mitochondria (Mit) and the rough endoplasmic reticulum (RER) are highlighted with blue and green arrows, respectively.

### Effects of ES on enzyme activities and synthetic functions in hepatocytes

Hepatocyte enzyme activities were measured. For ALT, AST, and LDH, we found an interaction between ES and time. ALT, AST, and LDH activities were significantly higher in the ES-24 h group compared with control-24 h, in the ES-48 h group compared with control-48 h, and in the ES-48 h group compared with ES-24 h. For ALP, GGT, and LAP, we found no interaction between ES and time. ALP, GGT, and LAP activities were increased after ES treatment compared with the control group (Fig. 6A).

**Fig 6.**
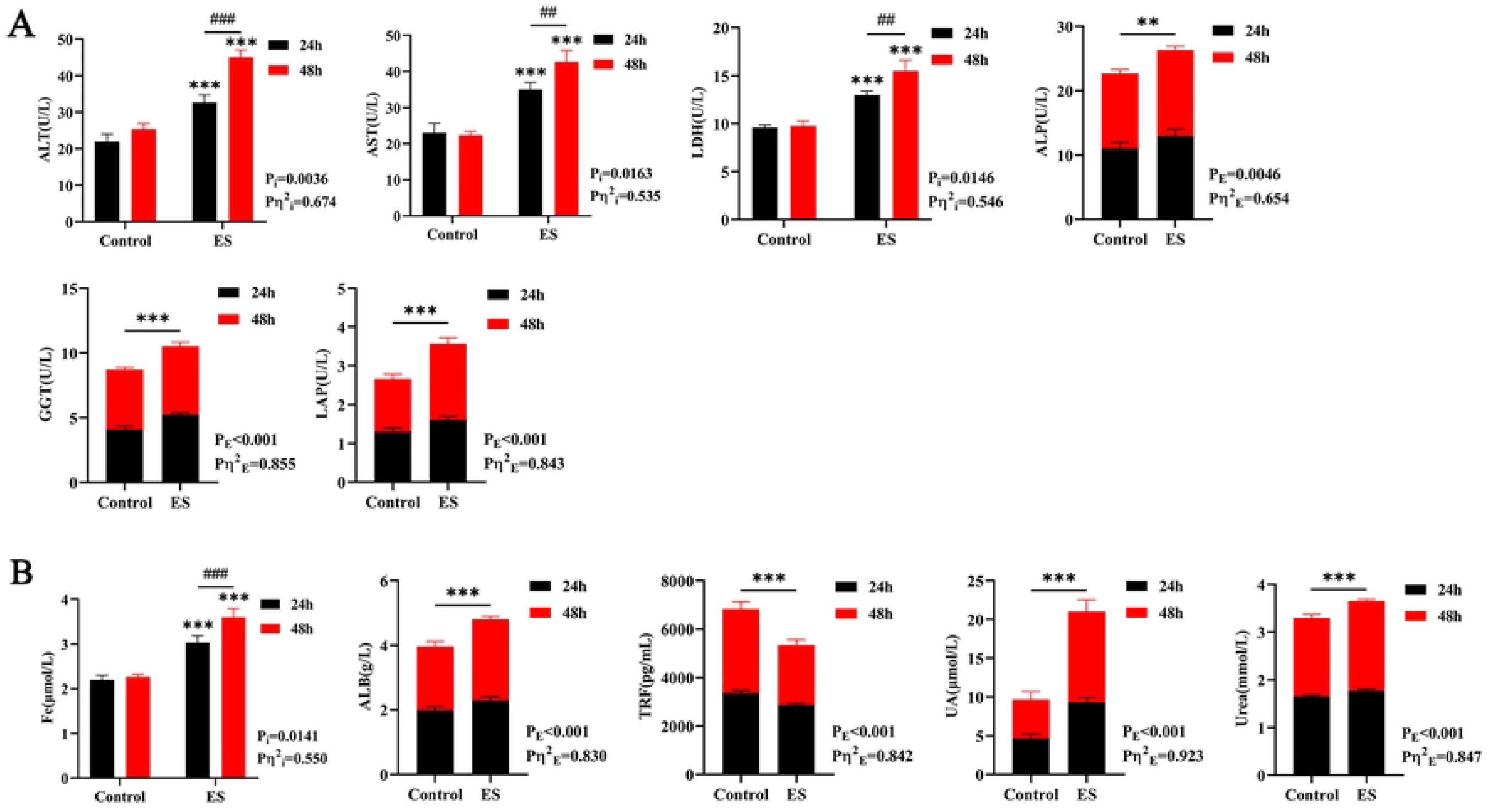
Hepatocyte enzyme activities and synthetic function indices after ES treatment. (A) Activity of six hepatocyte enzymes. (B) Levels of five hepatocyte synthetic substances. \**P* < 0.05 vs. control group, #*P* < 0.05.

Hepatocyte synthesis function was analyzed. For Fe, we found an interaction between ES and time. The Fe content in the ES-24 h group was higher than in control-24 h, it was higher in the ES-48 h group than in control-48 h, and it was higher in the ES-48 h group than in ES-24 h. For ALB, TRF, UA, and urea, we found no interaction between ES and time. ALB, UA, and urea levels were significantly higher and TRF levels were significantly lower after ES treatment compared with the control group (Fig. 6B).

### ES inhibits glucose metabolism in hepatocytes

Glu and LA levels in the culture medium can reflect changes in sugar metabolism (Fig. 7A). The Glu content was lower in the ES-48 h group than in control-48 h and ES-24 h, indicating that Glu was gradually consumed upon ES treatment. The content of LA was decreased after ES treatment, but it was not significantly different between the ES-24 h and ES-48 h groups, indicating that the production of LA was reduced after ES treatment.

**Fig 7.**
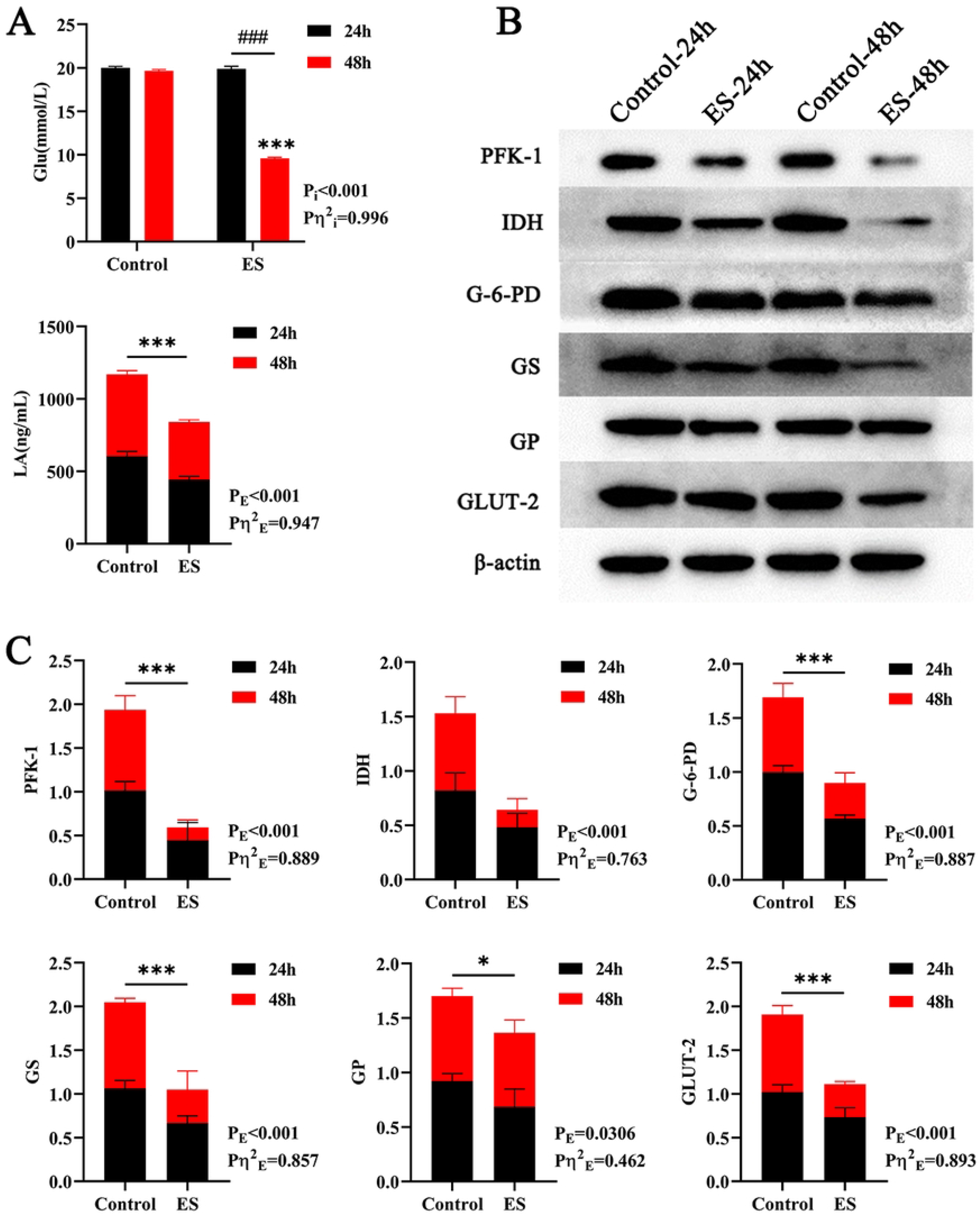
Effects of ES on glucose metabolism in hepatocytes. (A) The glucose and lactic acid contents reflect the cellular glucose metabolism level. (B) Western blot showed the expression levels of six key enzymes of glucose metabolism. (C) The relative expression levels of six proteins were obtained by ImageJ software. \**P* < 0.05 vs. control group, #*P* < 0.05.

PFK-1, IDH, G-6-PD, GS, GP, and GLUT-2 are the rate-limiting enzymes in multiple glucose metabolism pathways, and their expression levels can reflect the activity of cellular glucose metabolism. The expression levels of these enzymes were determined by WB (Fig. 7B). ImageJ software was used to obtain their relative expression levels (Fig. 7C). For all six enzymes, we found no interaction between ES and time. The protein expression levels of PFK-1, IDH, G-6-PD, GS, GP, and GLUT-2 were significantly lower in the ES groups than in the control groups, indicating that ES inhibits glucose metabolism in hepatocytes.

## Discussion

In contrast to previous studies using hydatid fluid (HF)[22], we used ES products from *E. granulosus* PSCs to characterize their effects on hepatocytes. The ES obtained was actually purified HF and did not require concentration[23]. ES isolation is equivalent to the removal of host impurities from HF and the reduction of its original state at the time of infection, because HF is mixed with host-related factors, and through our protocol, pure ES products can be obtained to improve the accuracy and reliability of the experiments. Although the number of PSCs was different in different hydatid cysts, the ES concentration was not the main research subject in our experiment, so in this experiment, we only used 100% ES. In terms of experimental design, different from the previously published randomized design[24], we adopted a factorial experimental design method, taking ES and time as two independent variables. In this way, we could not only observe the main effect of ES but also elucidate the interaction between ES and time, which greatly reduced the experimental error.

The CCK-8 results showed a trend of first increasing and then decreasing effects of ES on cell viability, which was different from the effects of microcyst fluid and hepatocytes[25]. Our results indicate that ES exerts its inhibitory effects on material consumption in hepatocytes within 48 h. In in vitro experiments, no fresh ES is produced, so after 48 h, the inhibitory effects of ES were weakened, and the proliferation activity of the cells was gradually enhanced. This situation is inevitable in in vitro experiments, so we chose 48 h as the maximum culture time. However, in animal experiments, this problem may be overcome to a certain extent.

During the natural development of CE, apoptosis may play a bifunctional role in the *host–Echinococcus* metabolite relationship[18]. This is consistent with our experimental results; both EdU and apoptosis experiments reflect the inhibitory and pro-apoptotic effects of ES on hepatocytes.

To the best of our knowledge, the structural changes of hepatocytes after hydatid cyst infection have not been characterized. Significant changes in hepatocyte structure were observed after ES treatment. Light microscopy revealed that hepatocyte morphology mainly changed from round or oval to polygonal or spindle shaped. SEM analysis revealed that the morphology of hepatocytes was extremely irregular, the cell structure collapsed, the cell membrane was broken, surface foaming and depression were prominent, and the microvilli fell off. TEM revealed that the nuclei were changed significantly, with some nuclei being pyknotic, the CH edges converged, and the NE surface was rough. The number and volume of Mit in the cytoplasm were significantly reduced, and in some cells, the RER was significantly expanded. The above changes became more obvious with the prolongation of time. These results suggest that ES may cause necrosis of host hepatocytes by disrupting the cell membrane, nucleus, Mit, and/or RER during infection, but the specific mechanism remains unclear.

The liver is the most enzyme-rich organ in the human body. ALT, ALP, GGT, and LAP in human serum are mainly derived from hepatocytes. ALT, ALP, and LAP are mainly found in the cytoplasm of hepatocytes, while GGT is mainly found in the cell membrane and microsomes of hepatocytes[26–29]. AST is mainly derived from cardiomyocytes, followed by hepatocytes, and most AST is located in mitochondria in hepatocytes[30]. LDH is a glycolytic enzyme mainly found in the myocardium, skeletal muscle, and kidney tissue, followed by the liver, and is mostly located in the cytoplasm in hepatocytes[31]. Only small amounts of the above six enzymes are released into the blood under normal conditions. When hepatocyte injury or necrosis occurs, the permeability of the cell membrane increases, which can result in increased levels of these enzymes in the blood. Therefore, ALT, AST, ALP, LDH, GGT, and LAP are commonly used indicators of hepatocyte injury in clinical practice[32–34]. In this study, ALT, AST, LDH, GGT, ALP, and LAP activities in the hepatocyte culture medium were increased after ES treatment compared with the control group, and the changes in ALT, AST, and LDH activities were time dependent. These results were consistent with the data obtained from the blood of CE patients[35]. The elevation of ALT and AST activities indicated that ES damaged not only the cell membrane but also mitochondria to a certain extent, which was consistent with our SEM and TEM observations. This may be one of the causes of hepatocyte necrosis in CE, but the specific mechanism needs to be further studied. The elevation of LDH, GGT, ALP, and LAP activities is also indicative of hepatocyte injury. These changes have certain clinical significance for the diagnosis of CE complicated with liver fibrosis, cirrhosis, and cholestatic jaundice, but these indices need to be analyzed together with bilirubin.

After ES treatment, the Fe content and the synthesis of ALB, UA, and urea in hepatocytes increased, while the synthesis of TRF decreased, and the Fe levels were time dependent. ALB, which is synthesized by the liver, plays an important role in maintaining blood colloid osmotic pressure, transport of metabolites, and nutrition in vivo. ALB content is an important indicator of liver synthesis function[36]. Increased ALB synthesis may be caused by decreased colloid osmolality, suggesting that the host body may suffer from a decreased plasma colloid osmolality in the early stage of hydatid infection and stimulate hepatocytes to synthesize ALB. PSCs may need hepatocyte ALB as a carrier to transport nutrients into the hydatid cavity to promote the growth and development of the hydatid cyst. Furthermore, during the growth of PSCs, PSCs may secrete ALB-like substances, which may increase the pseudo synthesis of ALB. The reasons for the increase in ALB synthesis analyzed above are based on the data of this experiment, but there is no relevant literature to support it, and the specific reasons need to be further explored.

Fe is an essential element for the formation of heme, which is necessary for the synthesis of hemoglobin and myoglobin and is also essential for the promotion of vitamin B metabolism. TRF is mainly synthesized in the liver and is a kind of globulin that can bind to Fe^3+^ and performs the function of transporting Fe[37]. The increase in Fe levels suggests acute hepatocyte injury and the release of intracellular stores of Fe, which has the same causes as the increase of ALT. Decreased TRF can also indicate hepatocyte injury, and its synthetic capacity is reduced. Iron-deficiency anemia is likely to occur in the presence of decreased TRF synthesis and Fe loss, which suggests that we need to pay attention to anemia indicators in CE and actively intervene when necessary.

Ammonia, which is highly toxic to the central nervous system, is one of the end products of amino acid metabolism[38]. The liver is the only organ that can relieve ammonia toxicity. Urea is generated through the ornithine cycle in the liver, and most urea is excreted by the kidneys. Increased urea synthesis could reflect the increased content of intracellular ammonia and enhanced protein catabolism, indicating ES enhances protein utilization. The increase of serum ammonia can also cause mitochondrial dysfunction[39]. As mentioned above, ES can induce different degrees of damage to hepatocytes. If the release of ammonia is greater than the compensatory function of hepatocytes, ammonia cannot be detoxified, which may eventually lead to the occurrence of hepatic encephalopathy, which is a life-threatening complication[40]. However, the occurrence of hepatic encephalopathy in CE patients has not been reported, which suggests that the PSCs use the host more than complete destruction. In addition, the increased protein demand of PSCs may be the main reason for the increase in ammonia levels, which may be considered to be consistent with the increase in ALB levels.

As a metabolite of purine in nuclear proteins and nucleic acids, UA is mainly synthesized in hepatocytes. UA levels reflect the filtration function of glomeruli and the reabsorption function of renal tubules and also serve as the main diagnostic basis for gout[41]. Increased UA synthesis reflects the enhanced utilization of nuclear proteins and nucleic acids in the liver by ES, which promotes the excessive breakdown of nucleic acids and may cause complications such as gout. Increased UA levels indicate that we need not only treat the primary disease but also to take measures to treat the corresponding complications. For example, oral antioxidant drugs can be used to prevent the oxidative decomposition of nucleic acids in cells, and when UA levels are increased, urine can be moderately alkalized with baking soda to promote UA excretion.

It is worth mentioning that almost none of the enzyme activity and synthetic function data obtained by us meet the diagnostic criteria in clinical practice. However, the aim of our experiment was to analyze the effects of ES on hepatocytes at the cellular level.

Glu is an essential nutrient for hepatocyte growth, providing cells with energy and a carbon source. The physiological significance of anaerobic oxidation of glucose lies in the rapid supply of energy without using oxygen. Our experiments showed that the Glu content in the culture medium was decreased after ES treatment, indicating that the utilization of Glu in hepatocytes was increased. Both LA content and PFK-1 expression were decreased, and the combination of the two indicated that the anaerobic oxidation capacity of glucose in hepatocytes was significantly inhibited, leading to a decrease in LA production. Although LDH is also involved in this reaction and its content was increased, the increase is mainly caused by cell membrane disruption and extravasation, so the content of LDH in the cytoplasm was decreased, which further reduces the synthesis of LA.

The tricarboxylic acid cycle, which takes place in mitochondria, is a common pathway and hub for the metabolism of three major nutrients[42]. As a key enzyme in the tricarboxylic acid cycle, the decreased expression of IDH also indicates that ES has an inhibitory effect on the aerobic oxidation of glucose metabolism in hepatocytes. This, in combination with the mitochondrial damage observed by TEM, further indicates that ES has a significant inhibitory effect on hepatic glucose metabolism. However, it is not known whether ES destroys mitochondria or inhibits enzyme activity.

Glu is decomposed through aerobic and anaerobic oxidation, and NADPH and phosphoribose are generated through the pentose phosphate pathway, which can provide hydrogen and carbon sources for various anabolic processes[43]. G-6-PD is a key enzyme in the pentose phosphate pathway, and its activity is mainly influenced by NADPH/NADP^+^ in physiological situations. However, in pathological conditions such as upon ES treatment, the expression of G-6-PD is decreased, indicating that the enzyme is inhibited by ES, suggesting the production of NADPH and phosphoribose through the pentose phosphate pathway is decreased. The decrease in NADPH may lead to decreased synthesis of amino acids and lipids, decreased hydroxylation, or hemolytic jaundice, commonly known as favism, due to the inability to maintain the reduced state of glutathione. GGT is also involved in the metabolism of glutathione, and increased activity of GGT indicates severe effusion, while decreased GGT content in the cytoplasm suggests a high possibility of hemolytic jaundice, although this has not been reported in patients with CE. A decrease in phosphoribose levels may lead to decreased nucleic acid biosynthesis. These changes indicate that ES not only inhibits the activity of G-6-PD but may also cause a series of complications.

Both GS and GP, which are key enzymes in the glycogen synthesis and decomposition pathways, respectively, act on α-1,4-glycosidic bonds. The simultaneous reduction in the activities of GS and GP indicates that ES damages the structures of the two enzymes. The specific mechanism is still unclear. GLUT-2 is a glucose transporter that is responsible for glucose uptake and insulin secretion in hepatocytes. Decreased GLUT-2 expression indicates that hepatocytes have a decreased ability to take up glucose, which directly affects glucose metabolism. The decreased expression of GLUT-2 combined with the previously detected decrease in the content of Glu in the cellular environment suggests that although the ability of hepatocytes to take up glucose was reduced, glucose could still enter the cells directly through simple diffusion due to the increased membrane permeability, rather than carrier-facilitated translocation[44].

It should be noted that for many of the complications analyzed above, clinical data are lacking. Nevertheless, more attention should be paid to a series of clinical symptoms caused by CE.

Glucose anaerobic oxidation, the tricarboxylic acid cycle, and the pentose phosphate pathway were inhibited in hepatocytes, indicating that *E. granulosus* has an inhibitory effect on hepatocyte glucose metabolism. *E. granulosus* contains a complete glycolytic pathway, tricarboxylic acid cycle, and pentose phosphate pathway, but lacks the ability to de novo synthesize pyrimidines, purines, and most amino acids (except alanine, aspartate, and glutamate), so it relies on the host to provide essential nutrients for its survival[45]. Thus, the relationship between PSCs or *E. granulosus* and its host is not only that of “killer” and “victim,” but more like that of “kidnapper” and “hostage.”

## Conclusions

Research on the pathogenesis, treatment, and prevention of CE is advancing. In the present study, *E. granulosus* ES products were isolated and their effects on hepatocytes were analyzed in vitro, to realistically simulate the interaction between PSCs and the surrounding hepatocytes. The inhibitory effects of ES on hepatocyte proliferation and the pro-apoptotic effects were verified experimentally. The structure of hepatocytes was damaged to a certain extent, the enzymatic activity of hepatocytes was increased, the synthesis function of hepatocytes was enhanced, and the biological effects of various glucose metabolism pathways were significantly inhibited after ES treatment. We analyzed the possible complications caused by various changes and advocated corresponding measures. We also proposed possible mechanisms by which hepatocyte necrosis may be induced in CE, but the specific mechanism requires further study.

## References

1. Thompson RC. Biology and Systematics of Echinococcus. Adv Parasitol. 2017;95:65–109. Doi:10.1016/bs.apar.2016.07.001. PMID: 28131366.

2. Flisser A. Eliminating cystic echinococcosis in the 21st century. Lancet Infect Dis. 2018;18(7):703–4. doi:10.1016/s1473-3099(18)30299-8. PMID: 29793822.

3. Fu MH, Wang X, Han S, Guan YY, Bergquist R, Wu WP. Advances in research on echinococcoses epidemiology in China. Acta Trop. 2021;219:105921. doi:10.1016/j.actatropica.2021.105921. PMID: 33878307.

4. Hsu TL, Lin G, Koizumi A, Brehm K, Hada N, Chuang PK, et al. The surface carbohydrates of the Echinococcus granulosus larva interact selectively with the rodent Kupffer cell receptor. Mol Biochem Parasitol. 2013;192(1-2):55–9. doi: 10.1016/j.molbiopara.2013.12.001. PMID: 24361107.

5. Nakao M, Lavikainen A, Yanagida T, Ito A. Phylogenetic systematics of the genus Echinococcus (Cestoda: Taeniidae). Int J Parasitol. 2013;43(12-13):1017–29. doi: 10.1016/j.ijpara.2013.06.002. PMID: 23872521.

6. Wan L, Wang T, Cheng L, Yu Q. Laparoscopic Treatment Strategies for Liver Echinococcosis. Infect Dis Ther. 2022;11(4):1415–26. doi: 10.1007/s40121-022-00664-2. PMID: 35751754.

7. Bastid C, Terraz S, Toso C, Chappuis F, Spahr L, Bresson-Hadni S. [Update on cystic echinococcosis of the liver]. Rev Med Suisse. 2021;17(748):1466–73. PMID: 34468098.

8. Wen-Jun Z, Xiu-Min H, Ya-Min G. [Progress in researches of benzimidazole in treatment of echinococcosis]. Zhongguo Xue Xi Chong Bing Fang Zhi Za Zhi. 2017;29(4):530–3. doi: 10.16250/j.32.1374.2017057. PMID: 29508601.

9. Wang S, Ma Y, Wang W, Dai Y, Sun H, Li J, et al. Status and prospect of novel treatment options toward alveolar and cystic echinococcosis. Acta Trop. 2022;226:106252. doi: 10.1016/j.actatropica.2021.106252. PMID: 34808118.

10. Brehm K, Koziol U. Echinococcus-Host Interactions at Cellular and Molecular Levels. Adv Parasitol. 2017;95:147–212. doi: 10.1016/bs.apar.2016.09.001. PMID: 28131363.

11. Virginio VG, Monteiro KM, Drumond F, de Carvalho MO, Vargas DM, Zaha A, et al. Excretory/secretory products from in vitro-cultured Echinococcus granulosus protoscoleces. Mol Biochem Parasitol. 2012;183(1):15–22. doi: 10.1016/j.molbiopara.2012.01.001. PMID: 22261090.

12. Jiménez M, Stoore C, Hidalgo C, Corrêa F, Hernández M, Benavides J, et al. Lymphocyte Populations in the Adventitial Layer of Hydatid Cysts in Cattle: Relationship With Cyst Fertility Status and Fasciola Hepatica Co-Infection. Vet Pathol. 2020;57(1):108–14. doi: 10.1177/0300985819875721. PMID: 31526120.

13. Spotin A, Majdi MM, Sankian M, Varasteh A. The study of apoptotic bifunctional effects in relationship between host and parasite in cystic echinococcosis: a new approach to suppression and survival of hydatid cyst. Parasitol Res. 2012;110(5):1979–84. doi: 10.1007/s00436-011-2726-4. PMID: 22167369.

14. Noori J, Spotin A, Ahmadpour E, Mahami-Oskouei M, Sadeghi-Bazargani H, Kazemi T, et al. The potential role of toll-like receptor 4 Asp299Gly polymorphism and its association with recurrent cystic echinococcosis in postoperative patients. Parasitol Res. 2018;117(6):1717–27. doi: 10.1007/s00436-018-5850-6. PMID: 29602972.

15. Pan W, Xu HW, Hao WT, Sun FF, Qin YF, Hao SS, et al. The excretory-secretory products of Echinococcus granulosus protoscoleces stimulated IL-10 production in B cells via TLR-2 signaling. BMC Immunol. 2018;19(1):29. doi: 10.1186/s12865-018-0267-7. PMID: 30355335.

16. Gottstein B, Soboslay P, Ortona E, Wang J, Siracusano A, Vuitton D. Immunology of Alveolar and Cystic Echinococcosis (AE and CE). Adv Parasitol. 2017;96:1–54. doi: 10.1016/bs.apar.2016.09.005. PMID: 28212788.

17. Soleymani N, Taran F, Nazemshirazi M, Naghibi A, Torabi M, Borji H, et al. Dysregulation of Ovine Toll-Like Receptors 2 and 4 Expression by Hydatid Cyst-Derived Antigens. Iran J Parasitol. 2021;16(2):219–28. doi: 10.18502/ijpa.v16i2.6271. PMID: 34557236.

18. Bakhtiar NM, Spotin A, Mahami-Oskouei M, Ahmadpour E, Rostami A. Recent advances on innate immune pathways related to host-parasite cross-talk in cystic and alveolar echinococcosis. Parasit Vectors. 2020;13(1):232. doi: 10.1186/s13071-020-04103-4. PMID: 32375891.

19. Atmaca HT. Determination of macrophage types by immunohistochemical methods in the local immune response to liver hydatid cysts in sheep. Acta Trop. 2022;229:106364. doi: 10.1016/j.actatropica.2022.106364. PMID: 35149039.

20. Yang HC, Xing ZK, Shao H, Tan XW, Wang EQ, Liao Y, et al. The expression of cytokeratin and apoptosis-related molecules in echinococcosis related liver injury. Mol Biochem Parasitol. 2022;248:111455. doi: 10.1016/j.molbiopara.2022.111455. PMID: 35016896.

21. Liu CS, Zhang HB, Yin JH, Jiang B, Han XM. Echinococcus granulosus: suitable in vitro protoscolices culture density. Biomed Environ Sci. 2013;26(11):912–5. doi: 10.3967/bes2013.020. PMID: 24331536.

22. Dos Santos GB, da Silva ED, Kitano ES, Battistella ME, Monteiro KM, de Lima JC, et al. Proteomic profiling of hydatid fluid from pulmonary cystic echinococcosis. Parasit Vectors. 2022;15(1):99. doi: 10.1186/s13071-022-05232-8. PMID: 35313982.

23. Wu J, Zhu Y, Zhou L, Lu Y, Feng T, Dai M, et al. Parasite-Derived Excretory-Secretory Products Alleviate Gut Microbiota Dysbiosis and Improve Cognitive Impairment Induced by a High-Fat Diet. Front Immunol. 2021;12:710513. doi: 10.3389/fimmu.2021.710513. PMID: 34745091.

24. Ma R, Qin W, Xie Y, Han Z, Li S, Jiang Y, et al. Dihydroartemisinin induces ER stress-dependent apoptosis of Echinococcus protoscoleces in vitro. Acta Biochim Biophys Sin (Shanghai). 2020;52(10):1140–7. doi: 10.1093/abbs/gmaa101. PMID: 33085744.

25. Liu C, Bi X, Fan H, Ma L, Ge RL. Microcyst fluid promotes the migration and invasion of fibroblasts in the adventitial layer of alveolar echinococcosis. Acta Trop. 2021;223:106084. doi: 10.1016/j.actatropica.2021.106084. PMID: 34389327.

26. Vujkovic M, Ramdas S, Lorenz KM, Guo X, Darlay R, Cordell HJ, et al. A multiancestry genome-wide association study of unexplained chronic ALT elevation as a proxy for nonalcoholic fatty liver disease with histological and radiological validation. Nat Genet. 2022;54(6):761–71. doi: 10.1038/s41588-022-01078-z. PMID: 35654975.

27. Wu Y, Zeng F, Sun L, Chen J, Wu S. ALP-activated probe for diagnosis of liver injury by multispectral optoacoustic tomography. Methods Enzymol. 2021;657:301–30. doi: 10.1016/bs.mie.2021.06.019. PMID: 34353492.

28. Liu T, Tian M, Wang J, Tian X, Liu J, Feng L, et al. Corrigendum to “Rational design of a fluorescent probe for the detection of LAP and its application in drug-induced liver injury” [Spectrochim. Acta Part A: Mol. Biomol. Spectrosc. 251 (2021) 119362]. Spectrochim Acta A Mol Biomol Spectrosc. 2021;261:120027. doi: 10.1016/j.saa.2021.120027. PMID: 34119766.

29. Zhang P, Jiang XF, Nie X, Huang Y, Zeng F, Xia X, et al. A two-photon fluorescent sensor revealing drug-induced liver injury via tracking γ-glutamyltranspeptidase (GGT) level in vivo. Biomaterials. 2016;80:46–56. doi: 10.1016/j.biomaterials.2015.11.047. PMID: 26706475.

30. Yao M, Wang L, You H, Wang J, Liao H, Yang D, et al. Serum GP73 combined AST and GGT reflects moderate to severe liver inflammation in chronic hepatitis B. Clin Chim Acta. 2019;493:92–7. doi: 10.1016/j.cca.2019.02.019. PMID: 30796901.

31. Lev-Cohain N, Sapir G, Harris T, Azar A, Gamliel A, Nardi-Schreiber A, et al. Real-time ALT and LDH activities determined in viable precision-cut mouse liver slices using hyperpolarized [1-(13) C]pyruvate-Implications for studies on biopsied liver tissues. NMR Biomed. 2019;32(2):e4043. doi: 10.1002/nbm.4043. PMID: 30575159.

32. Ahmed AE, Dahman B, Altamimi A, McClish DK, Al-Jahdali H. The aspartate aminotransferase/platelet count ratio index as a marker of dengue virus infection: Course of illness. J Infect Public Health. 2020;13(7):980–4. doi: 10.1016/j.jiph.2020.03.009. PMID: 32265161.

33. Hu X, Yang WX, Wang Y, Shao YX, Xiong SC, Li X. The prognostic value of De Ritis (AST/ALT) ratio in patients after surgery for urothelial carcinoma: a systematic review and meta-analysis. Cancer Cell Int. 2020;20:39. doi: 10.1186/s12935-020-1125-2. PMID: 32042266.

34. Qin C, Wei Y, Lyu X, Zhao B, Feng Y, Li T, et al. High aspartate aminotransferase to alanine aminotransferase ratio on admission as risk factor for poor prognosis in COVID-19 patients. Sci Rep. 2020;10(1):16496. doi: 10.1038/s41598-020-73575-2. PMID: 33020546.

35. Zhang J, Dong D, Yang J, Wang E, Chen X. Analysis of macrophage polarization and clinical characteristics in patients with hepatic cystic echinococcosis. J Trop Med. 2022;22(01):6–11.

36. Huang HY, Chen P, Liang XF, Wu XF, Wang CP, Gu X, et al. Dietary N-Carbamylglutamate (NCG) alleviates liver metabolic disease and hepatocyte apoptosis by suppressing ERK1/2-mTOR-S6K1 signal pathway via promoting endogenous arginine synthesis in Japanese seabass (Lateolabrax japonicus). Fish Shellfish Immunol. 2019;90:338–48. doi: 10.1016/j.fsi.2019.04.294. PMID: 31075404.

37. Xing G, Meng L, Cao S, Liu S, Wu J, Li Q, et al. PPARα alleviates iron overload-induced ferroptosis in mouse liver. EMBO Rep. 2022;23(8):e52280. doi: 10.15252/embr.202052280. PMID: 35703725.

38. Hadjihambi A, Cudalbu C, Pierzchala K, Simicic D, Donnelly C, Konstantinou C, et al. Abnormal brain oxygen homeostasis in an animal model of liver disease. JHEP Rep. 2022;4(8):100509. doi: 10.1016/j.jhepr.2022.100509. PMID: 35865351.

39. Angelova PR, Kerbert AJC, Habtesion A, Hall A, Abramov AY, Jalan R. Hyperammonaemia induces mitochondrial dysfunction and neuronal cell death. JHEP Rep. 2022;4(8):100510. doi: 10.1016/j.jhepr.2022.100510. PMID: 35845295.

40. Arjunan A, Sah DK, Jung YD, Song J. Hepatic Encephalopathy and Melatonin. Antioxidants (Basel). 2022;11(5). doi: 10.3390/antiox11050837. PMID: 35624703.

41. He J, Ye J, Sun Y, Feng S, Chen Y, Zhong B. The Additive Values of the Classification of Higher Serum Uric Acid Levels as a Diagnostic Criteria for Metabolic-Associated Fatty Liver Disease. Nutrients. 2022;14(17). doi: 10.3390/nu14173587. PMID: 36079844.

42. Rathod R, Gajera B, Nazir K, Wallenius J, Velagapudi V. Simultaneous Measurement of Tricarboxylic Acid Cycle Intermediates in Different Biological Matrices Using Liquid Chromatography-Tandem Mass Spectrometry; Quantitation and Comparison of TCA Cycle Intermediates in Human Serum, Plasma, Kasumi-1 Cell and Murine Liver Tissue. Metabolites. 2020;10(3). doi: 10.3390/metabo10030103. PMID: 32178322.

43. Zhang Z, TeSlaa T, Xu X, Zeng X, Yang L, Xing G, et al. Serine catabolism generates liver NADPH and supports hepatic lipogenesis. Nat Metab. 2021;3(12):1608–20. doi: 10.1038/s42255-021-00487-4. PMID: 34845393.

44. Chang YL, Chao AS, Chang SD, Cheng PJ. Placental glucose transporter 1 and 3 gene expression in Monochorionic twin pregnancies with selective fetal growth restriction. BMC Pregnancy Childbirth. 2021;21(1):260. doi: 10.1186/s12884-021-03744-2. PMID: 33773574.

45. Zheng H, Zhang W, Zhang L, Zhang Z, Li J, Lu G, et al. The genome of the hydatid tapeworm Echinococcus granulosus. Nat Genet. 2013;45(10):1168–75. doi: 10.1038/ng.2757. PMID: 24013640.

